# Replicability of multivariate brain-behaviour associations depends on clinical profile

**DOI:** 10.1101/2025.10.08.674881

**Authors:** Michelle Wang, Brent C. McPherson, Bratislav Misic, Franco Pestilli, Celia M. T. Greenwood, Jean-Baptiste Poline

## Abstract

Recent work suggests that thousands of individuals are required in multivariate brain-behaviour analyses to obtain consistently replicable results. Some believe, however, that smaller sample sizes may be sufficient if specific subpopulations are targeted. We investigate how sample size and cohort composition influence the replicability of Canonical Correlation Analysis (CCA) results using the UK Biobank (N=40,514). We apply CCA to diffusion-weighted magnetic resonance imaging (dMRI) phenotypes and cognitive assessment test scores. We define four participant cohorts based on clinical profile and find that, across all cohorts, sample sizes of _*≈*_500 are needed to obtain replicable canonical correlations and variable loadings. The most targeted cohort (comprising individuals with a history of psychoactive substance use) requires much fewer samples to achieve similar or greater correlations than the other cohorts. Our findings support the idea that moderate sample sizes from targeted cohorts can be sufficient for obtaining replicable brain-behaviour associations.

## Introduction

Traditional neuroimaging research has relied on small, locally acquired datasets that target specific scientific questions^1,2^: the median sample size of functional magnetic resonance imaging (fMRI) studies published in 2023 is estimated to be around 30^3^. This approach has resulted in unintentionally-biased sampling of research participants and has played a substantial part in the current replication crisis, with many studies lacking the power to be generalizable to new samples^4,5^. Acquiring neuroimaging data can be time-consuming and expensive, and it is often not realistic to expect individual neuroscientists to collect hundreds of samples in order to answer a single research question. As a solution, the field has embraced large-scale efforts to acquire and share large, deeply phenotyped datasets that maximize the availability of quality research data for the largest number of research questions^6–8^, thus enabling the use of more complex models.

For magnetic resonance imaging (MRI) neuroimaging research, notable large and accessible datasets include studies from the UK Biobank (UKBB; ∼40,000 participants)^9^, the Alzheimer’s Disease Neuroimaging Initiative (ADNI; ∼2500 participants)^10^, the Human Connectome Project (HCP; ∼1200 participants)^8^ and the Adolescent Brain and Cognitive Development study (ABCD; ∼12,000 participants)^11^. These datasets include not only neuroimaging information, but also data from other modalities such as rich behavioural assessments, clinical data and genetic information. The breadth of information collected can enable powerful multivariate analyses. One such multivariate method is Canonical Correlation Analysis (CCA)^12^, which can be thought of as a correlation between two (or more) domains. If variables in a dataset can be separated by modality – for example, into brain and behaviour features – then CCA finds linear combinations of variables in each modality such that these linear combinations (latent factors called “canonical axes”) are maximally correlated with each other. The interpretation of the features that matter most in this model is a descriptive way of showing the relationship between these features; variables with a stronger contribution to the axis are weighted higher, having a stronger “loading” onto the feature axis.

CCA has been used with samples like the HCP dataset to describe brain-behaviour relationships^13–15^. Unfortunately, CCA has notoriously poor replications to a single new sample^16,17^, meaning that variable loadings that are used to interpret the model can poorly reflect the true relationship of the underlying data. This limitation has recently been brought into the spotlight by Marek *et al*. (2022), who claimed that univariate and multivariate brain-wide association study (BWAS) methods like CCA need thousands of observations in order to accurately capture meaningful brain-behaviour relationships^18^. This has sparked many discussions in the neuroimaging community^19–37^, with some groups agreeing that such large sample sizes are indeed required^30,32^ and others arguing that smaller samples can be enough in some cases^23,24,26,31,36^.

Several of the responses to the Marek *et al*. (2022) study claim that more targeted samples (e.g., clinical populations) will create reproducible findings with only hundreds of samples due to the larger expected effect sizes in specific subpopulations^23,24,36^. This is an interesting distinction: most of the existing studies looking at how sample size impacts BWAS reproducibility used healthy populations with a large array of generic, non-specific features. Targeting more specific characteristics in a clinical sample may well improve the detection of relevant relationships, but this is yet untested.

In this study, we investigated the conjoint effects of sample size and sample composition on the reproducibility of multivariate models of brain-behaviour interactions in the UK Biobank (N=40,514). Unlike most previous work related to the reproducibility of brain-behaviour associations^18,24,30,31,33,35^, we used structural brain measures derived from diffusion magnetic resonance imaging (dMRI) instead of functional connectivity or grey matter features. Features based on dMRI metrics are currently underreported for this kind of multivariate analysis, especially considering its emerging utility in tracking disease progression in different dementias^38,39^. Using dMRI phenotypes and cognitive assessment test scores, we attempted to replicate previous findings from Marek *et al*. describing the impact of sample size on the replicability of CCA results and investigated whether subsetting the full sample into different “cohorts” influenced these results. In addition to comparing in-sample correlations with out-of-sample correlations, we also examined the set of variable contributions at different sample sizes. Overall, we find that CCA does not provide reliable results in samples with fewer than 500 individuals. However, we also show that individual clinical groups have a differing relationship than healthy controls. Our findings have important implications for smaller studies that are relying on more targeted populations to improve their effect sizes as an alternative to increasing sample size.

## Results

This study investigated the effect of cohort composition and sample size on brain-behaviour CCA reproducibility. We used data from 40,514 participants with brain imaging data from the UK Biobank dataset^9^. Brain data consisted of imaging-derived phenotypes (IDPs) from diffusion-weighted magnetic resonance imaging (dMRI). These included measures derived from diffusion tensor modelling (fractional anisotropy, FA; diffusion tensor mode, MO; and mean diffusivity, MD), as well as measures derived from Neurite Orientation Dispersion and Density Imaging (NODDI) (intracellular volume fraction, ICVF; isotropic or free water volume fraction, ISOVF; and orientation dispersion index, OD)^40^. The IDPs were obtained by computing the weighted mean of these dMRI measures over 27 white matter tracts^41^. Behavioural data consisted of key performance measures from nine cognitive function tests performed at the assessment centres. The tests targeted different domains of intelligence, including fluid intelligence/reasoning, matrix pattern completion, numeric memory, prospective memory, pairs matching, reaction time, symbol digit substitution, tower rearranging, and trail making. A total of 162 brain measures and 43 behavioural measures were included in this study.

From the Full cohort of 40,514 participants, we created three smaller cohorts targeting subsets of healthy participants (Healthy cohort, 6676 samples), participants with past psychoactive substance (e.g., alcohol, opioids, cannabinoids, sedatives or hypnotics, cocaine) use (Psychoactive cohort, 4725 samples), and participants with hypertension (Hypertension cohort, 7768 samples) (see **Methods** for details). Demographic information for each cohort is shown in Supplementary Table 1.

Subsamples of each cohort were generated at up to 30 logarithmically-spaced sample sizes, ranging from 50 to half of the total available data for each cohort. These subsamples were used to fit CCA models (Train set). For each model instance, a random half of the full cohort data (Test set) was used to assess the generalizability of the fitted CCA model on previously unseen data. We compared two types of CCA models: with and without cross-validation sampling. The model without cross-validation sampling consisted of a single CCA that was performed on the entire Train set, similar to many previous studies^13,15,18,31^. The model with cross-validation sampling was an ensemble model made up of 100 CCA instances, where each instance was fitted on a subset of the training data (see **Methods** for details); this is similar to the “repeated k-fold cross-validation” approach described in McPherson and Pestilli (2021)^14^, but has been adapted so that the model could be tested on fully held-out data. Each model was run 100 times for up to 30 sample sizes, yielding a total of up to 6,000 fitted model instances for each cohort.

For each fitted model instance, we obtained canonical correlations for the Train set and the Test set, as well as variable loadings (i.e., contributions) for each of the canonical axes for the Brain and Behaviour domains. Unless otherwise specified, results presented in this paper are for the first canonical axis (CA_1_).

### Replicable brain-behaviour CCA on the UK Biobank requires hundreds of samples

We investigated the reliability of brain-behaviour CCA at different sample sizes. As expected, for all four cohorts (Full, Healthy, Psychoactive, and Hypertension), mean canonical correlations for the Test set (*r*_*test*_) increase as the sample size increases, while those for the Train set (*r*_*train*_) decrease (Figure 1). The mean *r*_*test*_ at the largest sample size was 0.33 ± 0.01 for the Full cohort, 0.27 ± 0.01 for the Healthy cohort, 0.33 ± 0.01 for the Psychoactive cohort, and 0.30 ± for the Hypertension cohort. Across all cohorts, effect sizes were not statistically different from zero until at least a sample of N=213 and required a sample of at least N=487 to produce *r*_*test*_ values that are significantly higher than those produced by null permutation models. This indicates that sample sizes of at least hundreds are generally needed to produce reliable results in such an analysis. Similar results were observed with the CCA model without cross-validation sampling (Supplementary Figure 1).

**Figure 1:**
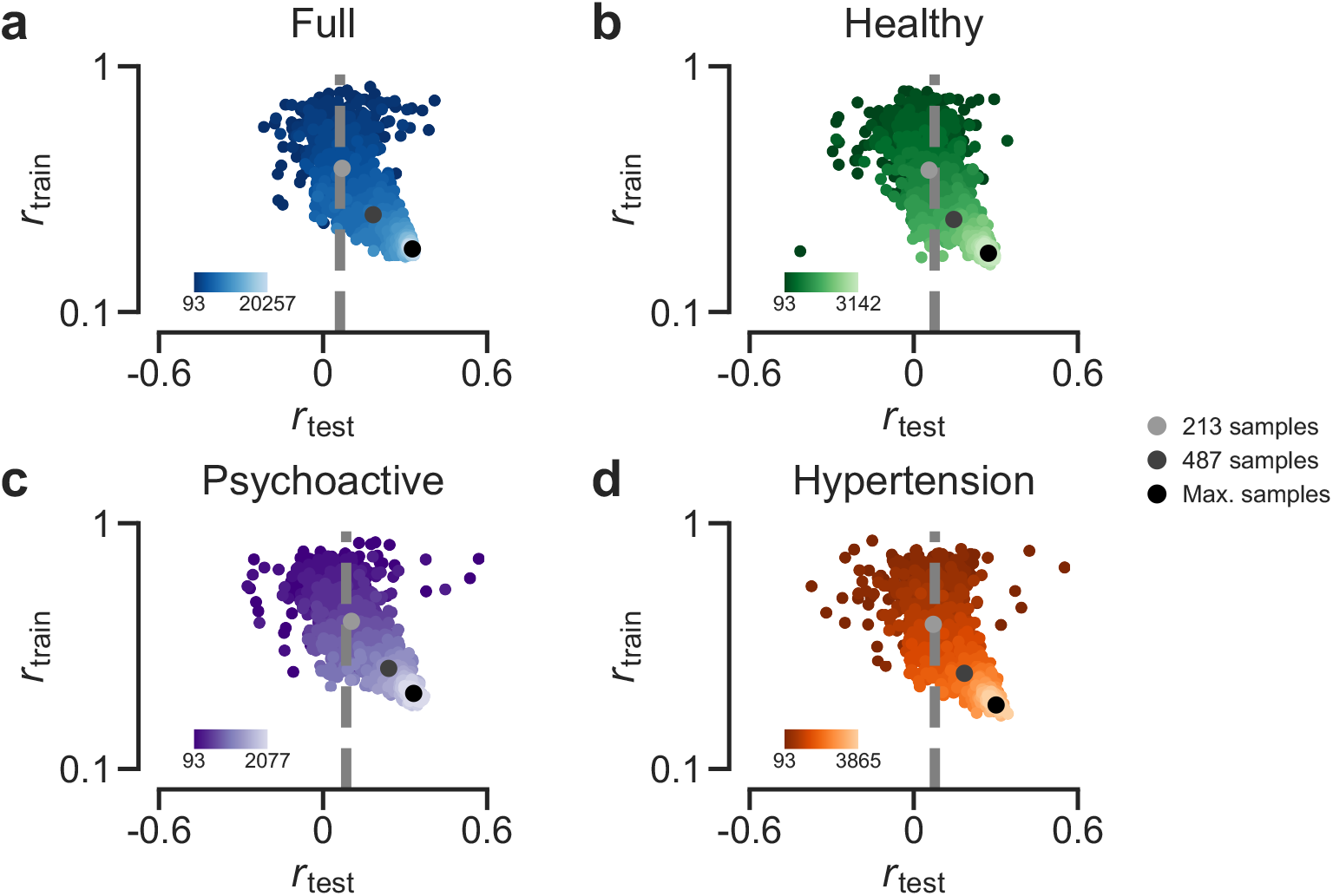
Train and test correlations across sample sizes and by cohort for the CCA model with cross-validation sampling. Train and test set correlations for each CCA model instance, coloured based on sample size. Grey dashed lines represent the 99th percentile of the null model distribution for the respective cohort (null models trained on largest available sample size). Larger dots represent mean *r*_*test*_ at specific sample sizes (light grey: 213, dark grey: 487, black: maximum). For visualization purposes, only sample sizes of at least 93 are shown (see Supplementary Figure 1 for all sample sizes). **a**: Full cohort. **b**: Healthy cohort. **c**: Psychoactive cohort. **d**: Hypertension cohort.

### Correlation strength on unseen data differs between cohorts

Cohorts targeting different clinical conditions within the sample had a moderating effect on the brain-behaviour relationship across sample sizes. Although all four cohorts showed mean Test set correlations (*r*_*test*_) that increased with sample size, the corresponding increase to the correlations differed by cohort. Notably, the Psychoactive cohort consistently showed higher correlations compared to all other three cohorts (Figure 2). At the sample size of 487, which corresponds to the lowest sample size for which there are sufficiently strong variable loadings for both Brain and Behavioural variables (as compared to null model loadings), the Psychoactive cohort had a mean *r*_*test*_ of 0.24 ± 0.05 (standard deviation), which is much higher than the mean *r*_*test*_ values for the Healthy cohort (0.15 ± 0.05) (Figure 2c). The Healthy cohort only reached a comparable *r*_*test*_ (0.24 ± 0.02) with a sample size of 1116, which is more than twice as large as what was needed for the Psychoactive cohort. Mean *r*_*test*_ values for the Hypertension and Full cohorts were close and largely remained in between the Healthy and Psychoactive cohort. Similar results were observed with the CCA model without cross-validation sampling (Supplementary Figure 2).

**Figure 2:**
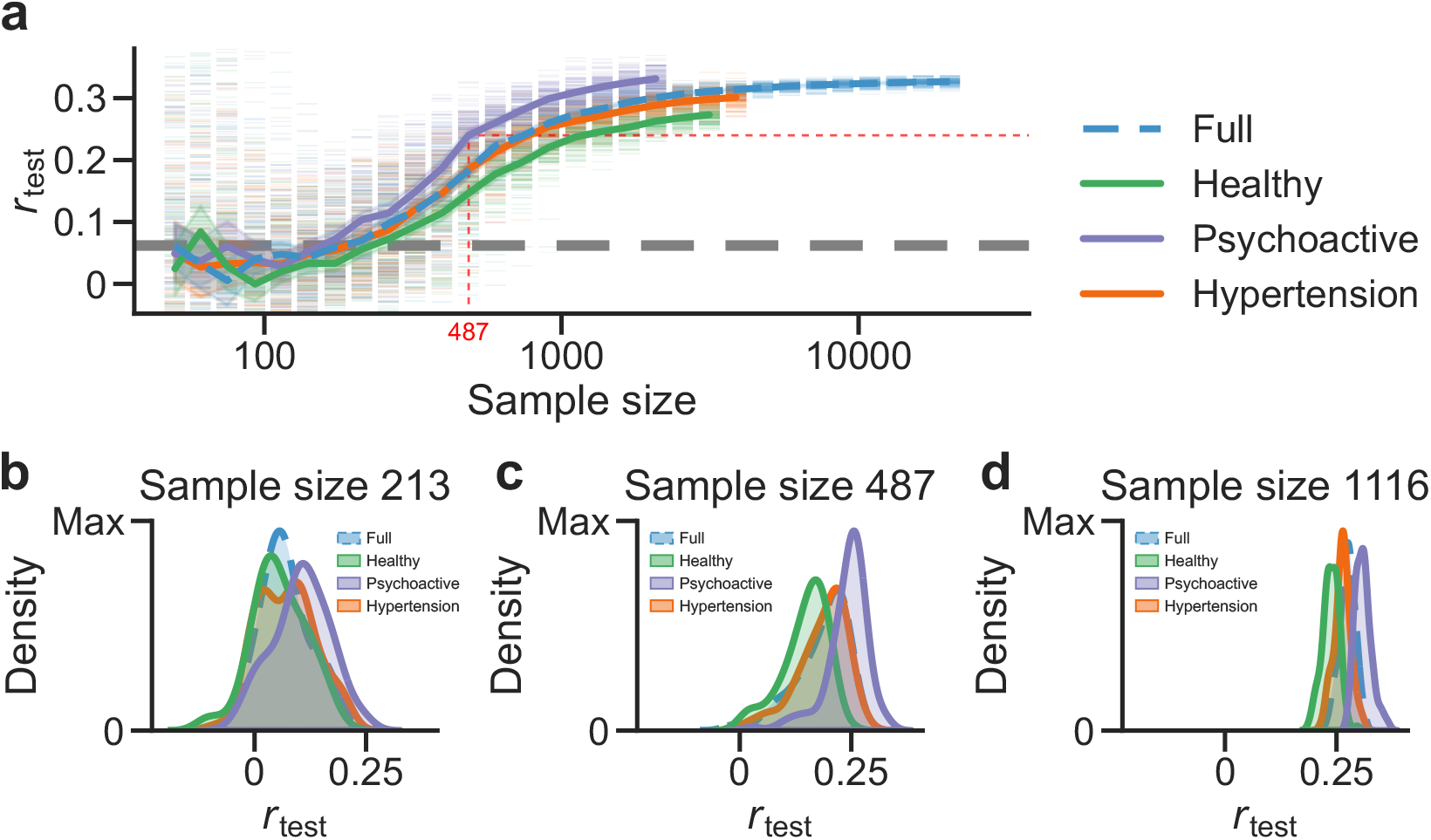
Replicability of canonical correlations across sample sizes and by cohort for the CCA model with cross-validation sampling. **a**: Average test set correlation by sample size for the Full cohort and for each other cohort featured in this analysis. The grey dashed line represents the 99th percentile of the null model distribution for the Full cohort at its greatest sample size. Error bands denote 95% confidence intervals. **b–d**: Kernel density estimation (KDE) distributions of the test set correlations for each cohort at sample sizes 213, 487, and 1116. The Psychoactive cohort consistently shows higher values compared to the other cohorts, while the Healthy cohort consistently shows lower values compared to the other cohorts.

To test the robustness of these results, we conducted an additional sensitivity analysis (Supplementary Figure 3) where we varied the number of features used as input to the CCA model (see **Methods**). We compared the minimum sample size needed for each cohort to reach a mean *r*_*test*_ of 0.24 (corresponding to the *r*_*test*_ at sample size 487 for the Psychoactive cohort in Figure 2). The Psychoactive cohort consistently required the lowest sample size (between 487 and 907) compared to the Full (between 907 and 1116), Healthy (between 1372 and 2077), and Hypertension (between 907 and 1372) cohorts (Supplementary Figure 4).

### Variable loading order is preserved across sample sizes

We compared the average variable loadings across the repeated models at each sample size for the Brain and Behaviour domains (Figure 3). For the Full cohort, the strongest positive and negative loadings (as determined by comparing with null model percentiles) are generally stable for sample sizes starting from around 500. This suggests that CCA with moderate sample sizes still provides a consistent estimation of the factor structure, even if the canonical correlations or factor loadings themselves are weaker. The variance of these estimates is relatively uniform for all variables at a given sample size and decreases as sample size increases (Supplementary Figure 5). For sample sizes lower than 100-200, the variable loadings are unstable: there is no clear pattern in the loading order, and the loadings have lower means as well as higher variances. Hence, CCA models fitted on these smaller sample sizes should not be interpreted since they are not replicable. All four cohorts show a similar loading order pattern, with some loadings passing the null model threshold at even lower sample sizes (e.g., Brain loadings for the Psychoactive cohort at N=262). Similar results were observed with the CCA model without cross-validation sampling (Supplementary Figure 6–7).

**Figure 3:**
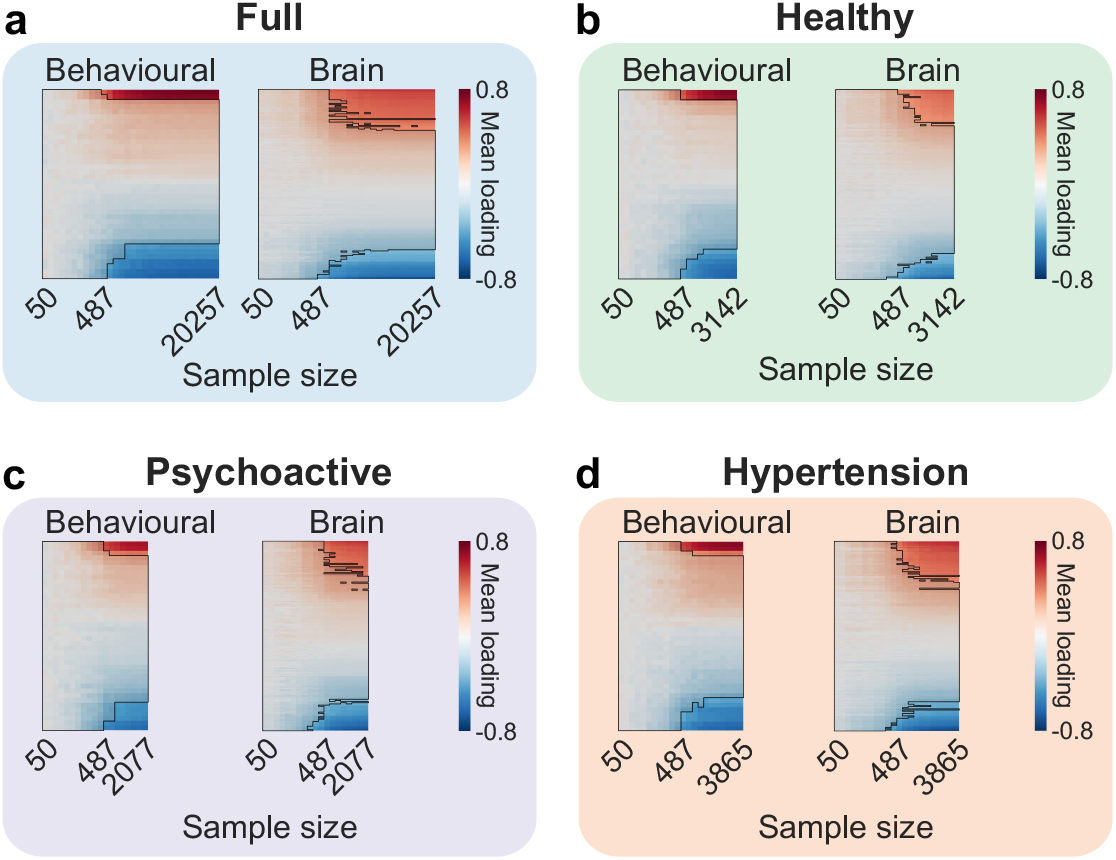
Mean loadings by sample size for brain and behavioural variables for each cohort for the CCA model with cross-validation sampling. Variable order is based on loading strengths at the largest sample size for each cohort. The saturation indicates the strength of the loading. Loadings whose mean value falls within the 2.5–97.5 null model percentile range are masked. **a**: Full cohort. **b**: Healthy cohort. **c**: Psychoactive cohort. **d**: Hypertension cohort.

Loadings that pass the null model threshold broadly agree between the four cohorts, though some minor differences exist (Supplementary Figures 8–11). Specifically, brain features related to the ICVF fraction of thalamo-cortical connections were the strongest positive loadings, and the MD of the same thalamo-cortical connections were the weakest, consistent with established relationships of anisotropic and isotropic diffusion features^42–44^. In general, the strongest positive behavioural loadings were related to high task performance, especially for the symbol digit substitution task, while the reaction time and trail-making tasks consistently had the strongest negative behavioural loadings.

### Cross-validation sampling reduces effect size inflation, but only at low sample sizes

Without cross-validation sampling, CCA models trained on small samples show severe overfitting. For the Full cohort, at the smallest sample size of 50, the mean *r*_*train*_ for the model without cross-validation sampling is 1.00 ± 0.00, while the mean *r*_*test*_ is −0.03 ± 0.34. In contrast, for the model with cross-validation sampling, the mean *r*_*train*_ and *r*_*test*_ are 0.01 ± 0.24 and −0.01 ± 0.20 respectively. With larger sample sizes, the two CCA models produce much closer results for both *r*_*train*_ and *r*_*test*_ (Supplementary Figure 12). We observe similar results for all four cohorts.

When it comes to the effect size inflation, as measured by the difference between Train set and Test set correlations, we find that CCA with cross-validation sampling reduces effect size inflation compared to CCA without cross-validation sampling, though only for sample sizes up to around 100 (Figure 4). This finding is not unexpected since a larger Train set already reduces chances of overfitting. However, much larger sample sizes (greater than N=1000) are needed to obtain an inflation value that is close to zero. Cross-validated and non cross-validated models both have the same levels of overfitting of sample sizes over sample sizes of 100.

**Figure 4:**
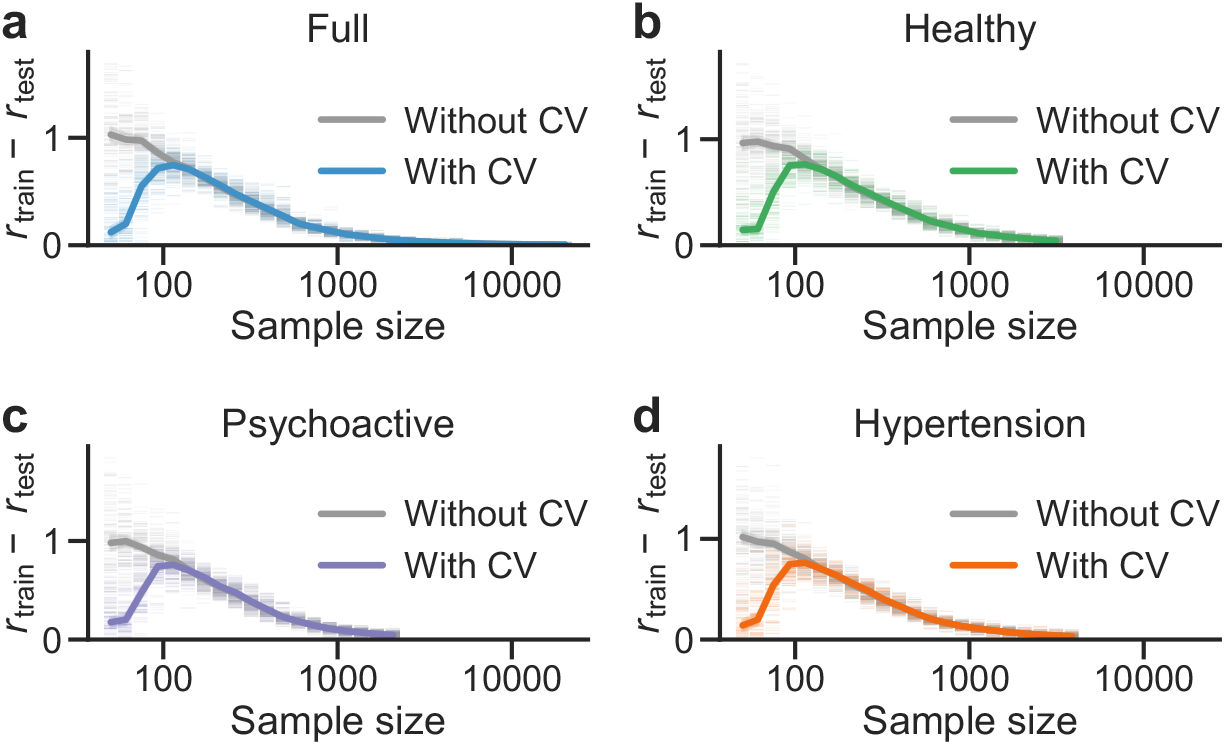
Comparison of effect size inflation between the two types of models for each cohort. Full lines: CCA model with cross-validation sampling. Dashed lines: CCA model without cross-validation sampling. Error bands denote 95% confidence intervals. **a**: Full cohort. **b**: Healthy cohort. **c**: Psychoactive cohort. **d**: Hypertension cohort.

## Discussion

In this study, we investigated the impact of cohort composition and sample size on the replicability of multivariate brain-behaviour relationships obtained from Canonical Correlation Analysis (CCA) methods. Utilizing data from the UK Biobank, we created training and test samples across a range of sample sizes from 50 to 20,257 for the full population and targeted clinical cohorts. We consistently reproduce general trends from Marek et al. 2022 showing that out-of-sample correlation increases as sample size increases. Sample sizes in the low hundreds or below led to non-replicable CCA model results and unstable variable loadings. However, although the largest sample sizes produce the best results, we find that sample sizes close to 500 are sufficient to obtain reproducible correlations and meaningful variable loadings with our specific study data. Moreover, variable loadings are stable even for these lower sample sizes (N_*≈*_500), and their order is preserved as sample size increases, indicating that they are reliably detecting a coherent underlying relationship of the variables. Our findings are therefore in line with several recent articles arguing that moderate sample sizes – in the order of hundreds, not thousands – may be enough for reproducible brain-behaviour associations^24,31^.

We found that using a targeted subset of participants with current or past psychoactive substance use resulted in higher replicability of brain-behaviour canonical correlations when compared to other participant cohorts at the same sample size. Psychoactive substance use is known to affect both cognitive function^45^ and white matter microstructure^46^: a change in either domain could change the observed relationship in our cohort^47^. The exact cause of this effect is unclear from our current analysis, and deeper interpretation of this specific finding was beyond the scope of our study. However, sensitivity analyses suggest that the behavioural domain could be important, as the effect is less marked with fewer behavioural measures (Supplementary Figure 3), but more systematic analyses would be needed to draw concrete conclusions.

Our main findings are consistent with the claim that lower sample sizes may be sufficient when effect sizes are expected to be larger due to study design^23,24,26,33,36^. It should be noted that our study only focused on varying cohort composition, while other factors such as feature selection likely also impact brain-behaviour associations. For example, collecting a more specialized set of features could lead to a sufficiently powered CCA with a much smaller sample. The same could be said about MRI modality, though a recent review of multivariate brain-behaviour relationship comparisons across clinical conditions found that cortical morphology and white matter association tracts had an overall stronger impact on a brain-behaviour axis than resting state fMRI^47^. In parallel, analytical choices such as using regularized CCA^48^ or other models (e.g., Bayesian methods) could impact the replicability of results as well. Such comparisons were beyond the scope of this study, but could form the basis of future work in this field.

We tested CCA models with and without cross-validation sampling (implemented as an ensemble model of 100 individual CCA models), and found that cross-validation sampling reduced the amount of overfitting on the training data, but only for very low sample sizes of around 100 or less. Even the largest sample size in this range was not enough to produce reliable brain-behaviour relationships in our other analyses. Nevertheless, using cross-validation sampling on small datasets can inform on sample size adequacy and give a more realistic expectation of model performance on previously unseen data.

In general, although our results support the notion that targeted cohort selection can reduce the sample size requirements for multivariate brain-behaviour analyses, we acknowledge that this may not always be the case depending on the cohort. Cohort specification is a procedure that can span multiple dimensions of subject characteristics: disease itself is a multidimensional trait, and it typically interacts with demographic, socioeconomic, and genetic features. It is likely that not all dimensions would impact brain-behaviour associations in the same manner. Therefore, in some cases it is possible that what appears to be a more clinically homogeneous cohort shows in practice smaller brain-behaviour effect sizes.

Overall, we believe this study answers some of the concerns regarding CCA analyses by directly testing the impact of sample size and composition on real data. We hope these findings can inform future studies when it comes to experimental design and data acquisition planning. In addition, we invite the scientific community to begin a discussion on the interpretation and downstream applications of CCA results. Specifically, what determines a significant model in the context of CCA has been poorly defined in previous work. In some cases, only results for the first canonical axis are presented^18,32^, although other axes may also produce significant canonical correlations. Moreover, the current literature seems to show little consideration for the reliability of feature contributions to canonical loadings (with some exceptions^32,35,48^). For example, in our study, we found that the first sample size with a significant first canonical axis was in fact lower than 487 for most cohorts; however, these canonical axes did not have strong loadings in both the Brain and Behavioural domains at these sample sizes. Would it be accurate to interpret these smaller sample sizes as adequate for capturing a multivariate comparison when none of the features provide statistically non-zero variable loadings? The statistical reliability of parameter estimates may be less relevant when the only interest is the descriptive ability of the canonical correlation between modalities. However, even in this context, often the strongest loading features of the axes are used to interpret the composition of the respective domains. A statistically derived threshold for parameters would allow for these and similar multivariate models to serve as testable, predictive models instead of purely descriptive.

To conclude, this study replicates some of the results found in Marek et al. (2022). Specifically, we find that out-of-sample CCA effect sizes are low at the sample sizes featured in most neuroimaging studies (≪500). Our findings complement previous work in the literature by showing that cohort composition can play a substantial role on the magnitude of these out-of-sample effect sizes. Further, when we looked beyond effect sizes, we found that we were able to obtain stable variable loadings at sample sizes of around 500. Overall, we find that reliable and meaningful CCA models still need large datasets, but with sample sizes on the order of hundreds, not thousands.

## Methods

### Dataset

This study used structural brain and behavioural measures from the UK Biobank (https://www.ukbiobank.ac.uk/; Application 45551). The UK Biobank project received approval from the National Information Governance Board for Health and Social Care and the National Health Service North West Centre for Research Ethics Committee (reference 11/NW/0382).

Informed consent was obtained from all participants, and all ethical regulations relevant to human research participants were followed.

We extracted dMRI measures and cognitive test results to form a Brain dataset and a Behavioural dataset respectively. Demographic, physical and physiological variables were treated as confounds and regressed out of the Brain and Behavioural datasets. Participants with completely missing data in any of the datasets were excluded from the entire analysis. Our final full dataset consisted of 40,514 participants.

UK Biobank participants typically underwent multiple visits (called “instances”) to the assessment centers. Unless otherwise specified in this text, data used in this study are from the first imaging visit (Instance 2). Data from subsequent imaging visits were not included because longitudinal features were beyond the scope of this analysis.. The UK Biobank data showcase website (https://biobank.ndph.ox.ac.uk/showcase/) was used to identify relevant categories and fields for the three datasets described below. Only numerical (integer, categorical, or continuous) data fields were considered.

### Brain variables

Structural brain variables consisted of pre-computed phenotypes derived from dMRI scans. We included a subset of the fields from Category 135 (“dMRI weighted means”), namely the weighted-mean measures for fractional anisotropy (FA), diffusion tensor mode (MO), mean diffusivity (MD), intracellular volume fraction (ICVF), isotropic or free water volume fraction (ISOVF), and orientation dispersion index (OD). These values are derived from fitted diffusion tensor and neurite orientation and dispersion density index (NODDI) model parameter values and were averaged within a set of 27 white matter tract regions. The dMRI acquisition methods and processing pipeline have been previously described by the UK Biobank neuroimaging analysis group^41^. A total of 162 structural brain variables were selected.

### Behavioural variables

A set of 43 cognitive function measures were selected for the Behavioural dataset. These measures consisted of results from a series of nine cognitive tests performed at the assessment centers: reaction time, numeric memory, fluid intelligence/reasoning, trail making, matrix pattern completion, tower rearranging, symbol digit substitution, prospective memory, and pairs matching (Category 100026: “Cognitive function”). For 15 measures, participants’ responses were binarized to indicate whether the participant answered correctly or not; these consisted of the fluid intelligence test items (Fields 4924, 4935, 4946, 4957, 4968, 4979, 4990, 5001, 5012, 5556, 5699, 5779, 5790, 5866) and Field 20018 “Prospective memory result”, where only participants who provided the right answer in their first attempt were considered correct. For cognitive assessments that involved multiple rounds (reaction time, numeric memory, trail making, matrix pattern completion, and pairs matching; Fields 398, 399, 400, 403, 404, 4255, 6333, 6770, 6771, 6772, 6773), individual round data were aggregated into summary measures of mean and standard deviation.

### Confounding variables

18 confounding variables were identified and regressed out of the Brain and Behavioural datasets. These measures were chosen based on methods described in previous work^13,14^. They consisted of the following:

1. Primary demographics, including sex and assessment center but excluding age (Category 1001, except Field 21003 “Age when attended assessment centre” and Field 34 “Year of birth”);
2. Blood pressure (Category 100011, except for Field 96 “Time since interview start at which blood pressure screen(s) shown”);
3. Handedness (Field 1707);
4. Pulse rate (Field 4194);
5. Standing height (Field 50);
6. Weight (Field 21002) and weight method (Field 21).

Some demographic fields, such as sex, were not recorded during the first imaging visit; in these cases, the value at Instance 0 was used for all participants.

Age-related data were not included as confound variables due to evidence that age may be more of an effect modifier than a confounder: brain-behaviour CCA associations are strongly linked with age, and including age as a confound reduces canonical correlations to near zero^14^.

### Cohort selection

Cohorts were formed based on the International Classification of Diseases 10 (ICD-10) summary diagnosis codes available for each participant (Fields 41270, 41202, 41204, and 41201). The selection criteria consisted of the following:

1. **Full**: all available participants (i.e., no filtering);
2. **Healthy**: participants without any diagnosis code;
3. **Psychoactive**: participants with code Z86.4 “Personal history of psychoactive substance abuse” or any code in the F10-F19 categories (“Mental and behavioural disorders due to psychoactive substance use”) (see https://icd.who.int/browse10/2019/en#/F10-F19 for more detailed information);
4. **Hypertension**: participants with code I10 “Essential (primary) hypertension”.

The final cohort sizes were the following: 40,514 for Full, 6676 for Healthy, 4725 for Psychoactive, and 7768 for Hypertension. By definition, the Full cohort contained all the participants in the other cohorts (as well as additional ones), and the Healthy cohort had no overlap with either the Psychoactive or the Hypertension cohort. Summary demographic information for each of the four cohorts is available in Supplementary Table 1. The Psychoactive and Hypertension cohorts had 1989 participants in common – these did not seem to be driving the effects we found, as removing them from the analysis still showed that the Psychoactive cohort needed less samples compared to the Healthy or Hypertension cohorts (Supplementary Figure 13).

### Data preprocessing

All categorical variables were one-hot encoded. Variables were removed if they had more than 50% missing data or if more than 80% of participants had the same value. The 80% threshold for low variability was chosen to ensure the impossibility of obtaining data with no variability at any point during the repeated 5-fold cross-validation procedure.

Preprocessing steps for confounding variables consisted of iterative imputation, inverse normal transformation and z-score standardization using the median and interquartile range. Squared versions of integer or continuous demographic variables were also added to account for some nonlinear effects when deconfounding.

Identical preprocessing pipelines were used for each of the Brain and Behavioural datasets. Variables were first normalized and z-scored (ignoring missing values). Then, a linear regression model was used to regress out the preprocessed confounding variables from the data. Principal Component Analysis (PCA) was then applied to reduce the dimension of the two datasets to 20 principal components (PCs) each. PCA was performed in a way that did not require imputing missing values: the data covariance matrix was first computed while ignoring missing data, then this (possibly non-diagonalizable) covariance matrix was projected to the nearest symmetric positive definite matrix, which was used in the eigenvector decomposition^13,14^.

### Canonical Correlation Analysis

Canonical Correlation Analysis (CCA) consists of solving the following optimization problem:

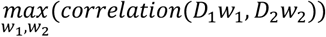

where *D*_1_ and *D*_2_ are two datasets (with observations as rows and features as columns), and *w*_1_ and *w*_2_ are weights that linearly combine columns from these datasets into *canonical factors* (or *canonical variables*) that are maximally correlated with each other. Additional pairs of canonical factors can be obtained by solving the same optimization problem, under the constraint that a pair of canonical factors cannot be correlated with another pair of canonical factors (i.e., they must be orthogonal). CCA can also be seen as finding a latent space such that projections of *D*_1_ and *D*_2_ onto that space have the highest correlation in the first latent dimension (first canonical axis, CA_1_), second highest correlation in the second latent dimension (CA_2_), and so on for all CAs.

In addition to the previously described canonical factors, the main results of an analysis involving CCA also include *canonical correlations* and *canonical loadings*. Canonical correlations are the correlations between each pair of canonical factors, and canonical loadings describe how features from the original data contribute to the canonical factors. Canonical loadings are obtained by correlating the original data matrices with the canonical factors.

### General data sampling and model fitting procedure

Data were split into Train and Test sets. The Train set was used for model fitting, including any cross-validation/ensembling. The Test set was used for model evaluation and always matched the maximum Train set size for each cohort (20,257 for the Full cohort, 3142 for the Healthy cohort, 2077 for the Psychoactive cohort, and 3865 for the Hypertension cohort). For each sample size, we generated a Train set by sampling (without replacement) from the participants who were not in the Test set. Sample sizes ranged from 50 to 20,257 (half of the total available data) and were chosen to be evenly spaced on a log scale. The exact sample sizes used were 50, 61, 75, 93, 114, 140, 173, 213, 262, 322, 396, 487, 599, 737, 907, 1116, 1372, 1688, 2077, 2555, 3142, 3865, 4755, 5848, 7194, 8849, 10884, 13388, 16468, and 20257. This process (including the initial 50% Test set sampling) was repeated 100 times for each cohort and each CCA model. All 30 sample sizes were used for the Full cohort; the first 21 sample sizes (up to 3142) were used for the Healthy cohort, the first 22 sample sizes (up to 3865) were used for the Hypertension cohort, and the first 19 sample sizes (up to 2077) were used for the Psychoactive cohort. Within a cohort, the same samples were used for the CCA model without cross-validation and the CCA model with repeated cross-validation.

To ensure that there was no data leakage between Train sets and Test sets, the parameters used in the preprocessing pipelines and the CCA model were estimated using data from the Train set only. Similarly, feature selection was done using only data from the Train set. Since each model fit was repeated 100 times on different Train sets, this resulted in potentially different numbers of features for the same model trained on different data, especially for smaller sample sizes.

### CCA without cross-validation sampling

We used the basic (non-regularized) CCA implementation from the CCA-Zoo Python package^50^ (version 1.10.11), which uses eigenvector decomposition to find the optimal transformation weights for the two datasets. The number of CAs was set to 20, which was the dimension of the Brain and Behavioural datasets after dimensionality reduction with PCA. This approach did not use cross-validation and consisted of training a single instance of the model on the entire Train set.

### CCA with cross-validation sampling

The cross-validation sampling approach was modified from the method described in a previous study by McPherson & Pestili^14^. Our approach consists of an ensemble model procedure where 5-fold cross-validation is repeated 20 times, yielding a total of 100 models. We then applied each fitted model to the full Train or Test set to obtain 100 sets of canonical factors. To account for arbitrary changes in direction that could have occurred between the 100 models, the canonical factors were aligned to the canonical factors from the CCA model without cross-validation sampling using the orthogonal Procrustes method^58–60^, using rotation matrices derived from the Train set canonical factors. The aligned canonical factors were then averaged to produce a single set of canonical factors for the overall (ensemble) model. Canonical correlations for the Train and Test sets were computed using the averaged canonical factors. Canonical loadings were obtained by correlating the normalized, z-scored and deconfounded data matrices (averaged from the 100 models) with the averaged canonical factors from the ensemble model.

### Null models

The null models were fitted in exactly the same way as the regular models, except that the Brain and Behavioural datasets were shuffled so that the participant order between them would not match. The maximum sample size was used for each of the four cohorts. Each null model was run 1000 times.

### Statistics and reproducibility

All analyses were performed using the Python programming language. The main packages used to implement the CCA models were scikit-learn^49^ and CCA-Zoo^50,51^. Other packages included pandas^52,53^ and NumPy^54^ for data wrangling, as well as Matplotlib^55,56^ and seaborn^57^ for plotting.

CCA models were run 100 times for each combination of sample size, cohort and model type (with or without cross-validation sampling). Null models were run 1000 times. Strong canonical loadings were defined as those whose mean values fell outside the 2.5–97.5 percentile range of the null model distributions.

## Data availability

Data used in this study can be obtained by applying for access to the UK Biobank database (https://www.ukbiobank.ac.uk/).

Source data for figures is available at https://github.com/neurodatascience/ukbb-cca and has been deposited to the Zenodo data repository^61^.

## Code availability

The Python source code for this project is publicly available at https://github.com/neurodatascience/ukbb-cca. The version of the code that produced results presented in the paper has been deposited to the Zenodo data repository^61^.

## Author Contributions

MW: Conceptualization, Data curation, Formal analysis, Funding acquisition, Methodology, Software, Visualization, Writing – original draft, Writing – review & editing.

BCM: Conceptualization, Methodology, Writing – original draft, Writing – review & editing. BM: Writing – review & editing.

FP: Writing – review & editing.

CG: Funding acquisition, Writing – review & editing.

JBP: Conceptualization, Funding acquisition, Methodology, Supervision, Writing – review & editing.

## Competing interests

The authors declare no competing interests.

## Acknowledgements

This research has been conducted using the UK Biobank Resource under Application Number 45551. This research was enabled in part by support provided by Calcul Québec and the Digital Research Alliance of Canada.

MW acknowledges support from the Canadian Institutes of Health Research (CIHR) (Canadian Graduate Scholarship – Doctoral 193412), the Brain Canada Foundation (Next Gen Award in Parkinson’s Disease Research), the Centre UNIQUE – Centre de recherche Neuro-IA du Québec (Doctoral Excellence Scholarship), the Fonds de Recherche du Québec – Santé (FRQS) (Bourse de doctorat en recherche BF2 329969), Parkinson Québec, and the Fonds de Recherche du Québec – Nature et technologie (FRQNT) (Bourse de maîtrise en recherche B1X 315843). The Brain Canada Next Gen Award in Parkinson’s Disease Research has been made possible by the Canada Brain Research Fund (CBRF), an innovative arrangement between the Government of Canada (through Health Canada) and Brain Canada Foundation, and by the Mireille and Steinberg Foundation and the Growling Beaver Brevet. The UNIQUE research centre is funded by the FRQNT Strategic Clusters Program.

BCM is part of the support staff of the Canadian Neuroanalytics Scholars (CNS) Program funded by The Hilary and Galen Weston Foundation and supported by Alberta Neuroscience, the Hotchkiss Brain Institute, The Neuro, and the Ontario Brain Institute.

CG was supported by the Healthy Brains, Healthy Lives initiative at McGill University, funded through support from the Canada First Research Excellence Fund (CFREF), Quebec’s Ministère de l’Économie, de l’Innovation et de l’Énergie (MEIE) and the Fonds de recherche du Québec (FRQS, FRQSC and FRQNT).

FP was supported by the following grants: Wellcome Trust 226486/Z/22/Z (Principal Investigator F. Pestilli); NINDS UM1NS132207, BRAIN CONNECTS: Center for Mesoscale Connectomics (Principal Investigator K. Ugurbil); and NINDS U24NS140384, BRAIN CONNECTS: The Axonal Projectome EXchange (APEX) (Principal Investigator F. Pestilli). FP thanks the Amazon Web Services Open Data Sponsorship Program for supporting data storage for brainlife.io.

JBP acknowledges support from the National Institutes of Health (NIH; NIH-NIBIB P41 EB019936 [ReproNim], NIH-NIMH R01 MH083320 [CANDIShare], NIH RF1 MH120021 [NIDM], and NIH-NIMH R01 MH096906 [Neurosynth]), the Canadian Institutes of Health Research (CIHR; PJT-185948), the Michael J. Fox Foundation, the Quebec Parkinson Network, the McConnell Brain Imaging Centre, the Canada First Research Excellence Fund, awarded to McGill University for the Healthy Brains for Healthy Lives initiative (NeuroHub), the Chan Zuckerberg Initiative (EOSS5-0000000401) and the Brain Canada Foundation with support from Health Canada through the Canada Brain Research Fund in partnership with the Montreal Neurological Institute.

## Notes

https://github.com/neurodatascience/ukbb-cca

https://www.ukbiobank.ac.uk/

### Competing Interest Statement

The authors have declared no competing interest.

### Summary of Updates

Minor revisions based on final reviewer comments and editor checklist.

## References

1. Button, K. S. et al. Power failure: why small sample size undermines the reliability of neuroscience. Nat. Rev. Neurosci. 14, 365–376 (2013).

2. Szucs, D. & Ioannidis, J. P. A. Empirical assessment of published effect sizes and power in the recent cognitive neuroscience and psychology literature. PLOS Biol. 15, e2000797 (2017).

3. Dockès, J., Oudyk, K., Torabi, M., Vega, A. I. de la & Poline, J.-B. Mining the neuroimaging literature. eLife 13, (2024).

4. Falk, E. B. et al. What is a representative brain? Neuroscience meets population science. Proc. Natl. Acad. Sci. 110, 17615–17622 (2013).

5. Bzdok, D. & Yeo, B. T. T. Inference in the age of big data: Future perspectives on neuroscience. NeuroImage 155, 549–564 (2017).

6. Bycroft, C. et al. The UK Biobank resource with deep phenotyping and genomic data. Nature 562, 203–209 (2018).

7. Thompson, P. M. et al. The ENIGMA Consortium: large-scale collaborative analyses of neuroimaging and genetic data. Brain Imaging Behav. 8, 153–182 (2014).

8. Van Essen, D. C. et al. The WU-Minn Human Connectome Project: An overview. NeuroImage 80, 62–79 (2013).

9. Miller, K. L. et al. Multimodal population brain imaging in the UK Biobank prospective epidemiological study. Nat. Neurosci. 19, 1523–1536 (2016).

10. Jack Jr., C. R. et al. The Alzheimer’s disease neuroimaging initiative (ADNI): MRI methods. J. Magn. Reson. Imaging 27, 685–691 (2008).

11. Casey, B. J. et al. The Adolescent Brain Cognitive Development (ABCD) study: Imaging acquisition across 21 sites. Dev. Cogn. Neurosci. 32, 43–54 (2018).

12. Hotelling, H. Relations Between Two Sets of Variates. Biometrika 28, 321–377 (1936).

13. Smith, S. M. et al. A positive-negative mode of population covariation links brain connectivity, demographics and behavior. Nat. Neurosci. 18, 1565–1567 (2015).

14. McPherson, B. C. & Pestilli, F. A single mode of population covariation associates brain networks structure and behavior and predicts individual subjects’ age. Commun. Biol. 4, 1–16 (2021).

15. Goyal, N., Moraczewski, D., Bandettini, P. A., Finn, E. S. & Thomas, A. G. The positive– negative mode link between brain connectivity, demographics and behaviour: a pre-registered replication of Smith et al. (2015). R. Soc. Open Sci. 9, 201090 (2022).

16. Wang, H.-T. et al. Finding the needle in a high-dimensional haystack: Canonical correlation analysis for neuroscientists. NeuroImage 216, 116745 (2020).

17. Zhuang, X., Yang, Z. & Cordes, D. A technical review of canonical correlation analysis for neuroscience applications. Hum. Brain Mapp. 41, 3807–3833 (2020).

18. Marek, S. et al. Reproducible brain-wide association studies require thousands of individuals. Nature 603, 654–660 (2022).

19. Cognitive neuroscience at the crossroads. Nature 608, 647–647 (2022).

20. Revisiting doubt in neuroimaging research. Nat. Neurosci. 25, 833–834 (2022).

21. Bandettini, P. A. et al. The challenge of BWAs: Unknown unknowns in feature space and variance. Med 3, 526–531 (2022).

22. Chakravarty, M. M. Controversies on brain-wide association studies: commentaries from the field. Aperture Neuro BWAS Editorials, 1–1 (2022).

23. Gratton, C., Nelson, S. M. & Gordon, E. M. Brain-behavior correlations: Two paths toward reliability. Neuron 110, 1446–1449 (2022).

24. Libedinsky, I. et al. Reproducibility of neuroimaging studies of brain disorders with hundreds-not thousands-of participants. 2022.07.05.498443 Preprint at 10.1101/2022.07.05.498443 (2022).

25. Kong, X.-Z., Zhang, C., Liu, Y. & Pu, Y. Scanning reproducible brain-wide associations: sample size is all you need? Psychoradiology 2, 67–68 (2022).

26. Rosenberg, M. D. & Finn, E. S. How to establish robust brain–behavior relationships without thousands of individuals. Nat. Neurosci. 25, 835–837 (2022).

27. Tiego, J. & Fornito, A. Putting behaviour back into brain–behaviour correlation analyses. Aperture Neuro BWAS Editorials, 1–4 (2022).

28. Uddin, L. Q. Brain–behavior associations depend heavily on user-defined criteria. Aperture Neuro BWAS Editorials, 1–2 (2022).

29. Valk, S. L. & Hettwer, M. D. Commentary on ‘Reproducible brain-wide association studies require thousands of individuals’. Aperture Neuro BWAS Editorials, 1–2 (2022).

30. Liu, S., Abdellaoui, A., Verweij, K. J. H. & van Wingen, G. A. Replicable brain–phenotype associations require large-scale neuroimaging data. Nat. Hum. Behav. 7, 1344–1356 (2023).

31. Spisak, T., Bingel, U. & Wager, T. D. Multivariate BWAS can be replicable with moderate sample sizes. Nature 615, E4–E7 (2023).

32. Helmer, M. et al. On the stability of canonical correlation analysis and partial least squares with application to brain-behavior associations. Commun. Biol. 7, 1–15 (2024).

33. Kang, K. et al. Study design features increase replicability in brain-wide association studies. Nature 636, 719–727 (2024).

34. Makowski, C. et al. Leveraging the adolescent brain cognitive development study to improve behavioral prediction from neuroimaging in smaller replication samples. Cereb. Cortex 34, bhae223 (2024).

35. Nakua, H. et al. Comparing the stability and reproducibility of brain-behavior relationships found using canonical correlation analysis and partial least squares within the ABCD sample. Netw. Neurosci. 8, 576–596 (2024).

36. DeYoung, C. G. et al. Beyond Increasing Sample Sizes: Optimizing Effect Sizes in Neuroimaging Research on Individual Differences. J. Cogn. Neurosci. 37, 1023–1034 (2025).

37. Lee, H. J., Dworetsky, A., Labora, N. & Gratton, C. Using precision approaches to improve brain-behavior prediction. Trends Cogn. Sci. 29, 170–183 (2025).

38. Huang, C.-C. et al. Diffusion and structural MRI as potential biomarkers in people with Parkinson’s disease and cognitive impairment. Eur. Radiol. 34, 126–135 (2024).

39. Spotorno, N. et al. Diffusion MRI tracks cortical microstructural changes during the early stages of Alzheimer’s disease. Brain 147, 961–969 (2024).

40. Zhang, H., Schneider, T., Wheeler-Kingshott, C. A. & Alexander, D. C. NODDI: practical in vivo neurite orientation dispersion and density imaging of the human brain. NeuroImage 61, 1000–1016 (2012).

41. Alfaro-Almagro, F. et al. Image processing and Quality Control for the first 10,000 brain imaging datasets from UK Biobank. NeuroImage 166, 400–424 (2018).

42. Lawrence, K. E. et al. Age and sex effects on advanced white matter microstructure measures in 15,628 older adults: A UK biobank study. Brain Imaging Behav. 15, 2813–2823 (2021).

43. Greenman, D. & Bennett, I. J. Aging of gray matter microstructure: A brain-wide characterization of age group differences using NODDI. Neurobiol. Aging 149, 34–43 (2025).

44. Hughes, E. J. et al. Regional changes in thalamic shape and volume with increasing age. NeuroImage 63, 1134–1142 (2012).

45. Ramey, T. & Regier, P. S. Cognitive Impairment in Substance Use Disorders. CNS Spectr. 24, 102–113 (2019).

46. Hampton, W. H., Hanik, I. M. & Olson, I. R. Substance Abuse and White Matter: Findings, Limitations, and Future of Diffusion Tensor Imaging Research. Drug Alcohol Depend. 197, 288–298 (2019).

47. Vieira, S. et al. Multivariate brain-behaviour associations in psychiatric disorders. Transl. Psychiatry 14, 231 (2024).

48. Mihalik, A. et al. Canonical Correlation Analysis and Partial Least Squares for Identifying Brain–Behavior Associations: A Tutorial and a Comparative Study. Biol. Psychiatry Cogn. Neurosci. Neuroimaging 7, 1055–1067 (2022).

49. Pedregosa, F. et al. Scikit-learn: Machine Learning in Python. J. Mach. Learn. Res. 12, 2825–2830 (2011).

50. Chapman, J. & Wang, H.-T. CCA-Zoo: A collection of Regularized, Deep Learning based, Kernel, and Probabilistic CCA methods in a scikit-learn style framework. J. Open Source Softw. 6, 3823 (2021).

51. Chapman, J., Wang, H.-T., Wells, L. & Wiesner, J. CCA-Zoo. Zenodo 10.5281/zenodo.5930463 (2022).

52. McKinney, W. Data Structures for Statistical Computing in Python. in 56–61 (Austin, Texas, 2010). doi:10.25080/Majora-92bf1922-00a.

53. Reback, J. et al. pandas-dev/pandas: Pandas 1.4.0. Zenodo 10.5281/zenodo.5893288 (2022).

54. Harris, C. R. et al. Array programming with NumPy. Nature 585, 357–362 (2020).

55. Hunter, J. D. Matplotlib: A 2D Graphics Environment. Comput. Sci. Eng. 9, 90–95 (2007).

56. Caswell, T. A. et al. matplotlib/matplotlib: REL: v3.5.1. Zenodo 10.5281/zenodo.5773480 (2021).

57. Waskom, M. L. seaborn: statistical data visualization. J. Open Source Softw. 6, 3021 (2021).

58. Milan, L. & Whittaker, J. Application of the Parametric Bootstrap to Models that Incorporate a Singular Value Decomposition. J. R. Stat. Soc. Ser. C Appl. Stat. 44, 31–49 (1995).

59. Petrican, R. & Fornito, A. Adolescent neurodevelopment and psychopathology: The interplay between adversity exposure and genetic risk for accelerated brain ageing. Dev. Cogn. Neurosci. 60, 101229 (2023).

60. Schönemann, P. H. A generalized solution of the orthogonal procrustes problem. Psychometrika 31, 1–10 (1966).

61. Wang, M. neurodatascience/ukbb-cca: 1.0.0. Zenodo 10.5281/zenodo.20026935 (2026).

