## Supplementary information for "Replicability of multivariate brain-behaviour associations depends on clinical profile"

| | All | Psychoactive | Healthy | Hypertension |
|---|---|---|---|---|
| N | 40,514 | 4725 | 6676 | 7768 |
| Age (years) | 64.2 ± 7.8 | 64.2 ± 7.8 | 64.2 ± 7.7 | 64.2 ± 7.8 |
| Weight (kg) | 76.3 ± 15.3 | 76.3 ± 15.4 | 76.5 ± 15.1 | 76.1 ± 15.5 |
| Sex | | | | |
| *Male* | 19,588 (48.3%) | 2267 (48.0%) | 3298 (49.4%) | 3710 (47.8%) |
| *Female* | 20,926 (51.7%) | 2458 (52.0%) | 3378 (50.6%) | 4058 (52.2%) |
| Ethnicity | | | | |
| *White* | 39,202 (96.8%) | 4580 (96.9%) | 6456 (96.7%) | 7505 (96.6%) |
| *Mixed* | 185 (0.5%) | 28 (0.6%) | 30 (0.4%) | 36 (0.5%) |
| *Asian/Asian British* | 428 (1.1%) | 49 (1.0%) | 74 (1.1%) | 77 (1.0%) |
| *Black/Black British* | 270 (0.7%) | 26 (0.6%) | 47 (0.7%) | 57 (0.7%) |
| *Chinese* | 118 (0.3%) | 7 (0.1%) | 19 (0.3%) | 29 (0.4%) |
| *Other* | 202 (0.5%) | 26 (0.6%) | 31 (0.5%) | 47 (0.6%) |
| Handedness | | | | |
| *Right* | 36,043 (89.0%) | 4187 (88.6%) | 5972 (89.5%) | 6914 (89.0%) |
| *Left* | 3807 (9.4%) | 470 (9.9%) | 602 (9.0%) | 732 (9.4%) |
| *Both* | 645 (1.6%) | 67 (1.4%) | 98 (1.5%) | 120 (1.5%) |

**Supplementary Table 1: Participant demographics for each cohort.** Age and weight are given as mean ± standard deviation.

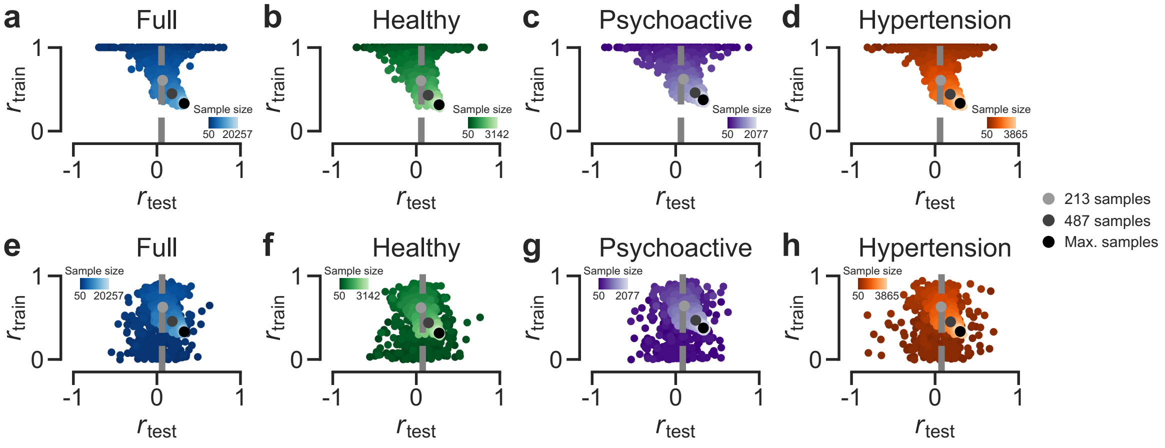


**Supplementary Figure 1: Train and test correlations across sample sizes and by cohort (CCA models <u>without</u> and <u>with</u> cross-validation sampling)**.
Train and test set correlations for each CCA model instance, coloured based on sample size. Grey dashed lines represent the 99th percentile of the null model distribution for the respective cohort (null models trained on largest available sample size). Larger dots represent mean $r_{test}$ at specific sample sizes (light grey: 213, dark grey: 487, black: maximum). **a–d**: CCA model without cross-validation sampling. **e–h**: CCA model with cross-validation sampling (all sample sizes).

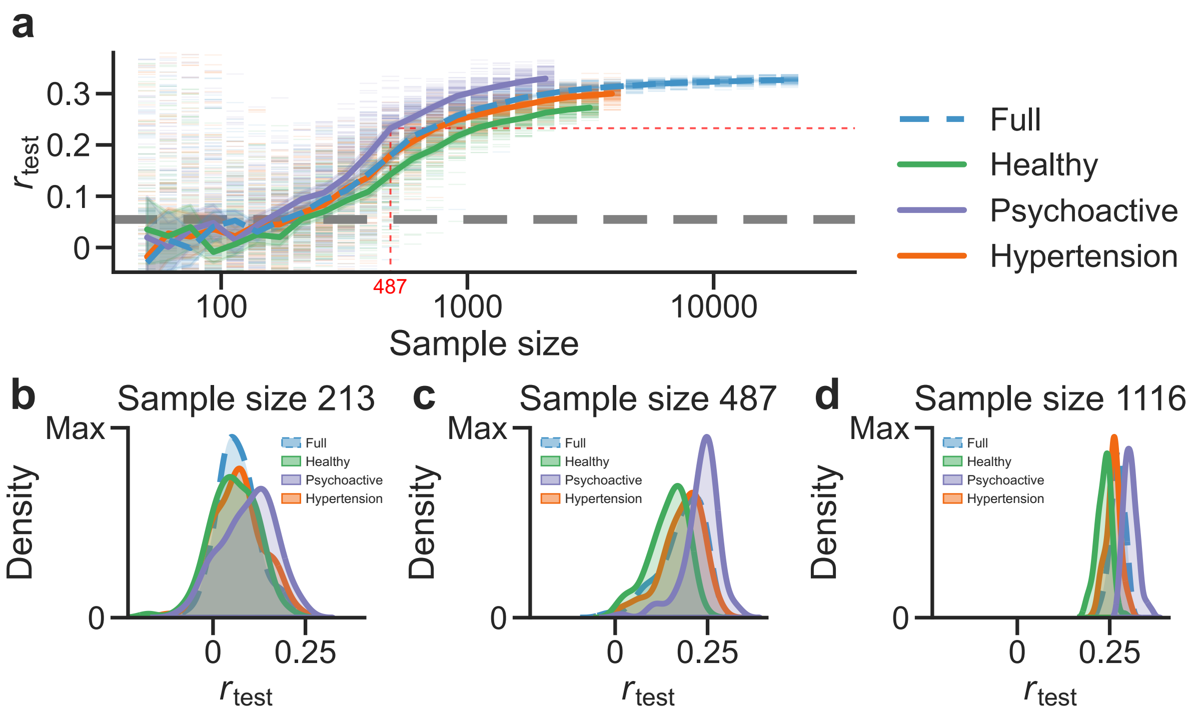


**Supplementary Figure 2: Replicability of canonical correlations across sample sizes and by cohort (CCA model without cross-validation sampling)**.
**a**: Average test set correlation by sample size for the Full cohort and for each other cohort featured in this analysis. Grey dashed line represents the 99th percentile of the null model distribution for the Full cohort at its greatest sample size. Error bands denote 95% confidence intervals. **b–d**: Kernel density estimation (KDE) distributions of the test set correlations for each cohort at sample sizes 213, 487, and 1116. The Psychoactive cohort consistently shows higher values compared to the other cohorts, while the Healthy cohort consistently shows lower values compared to the other cohorts.

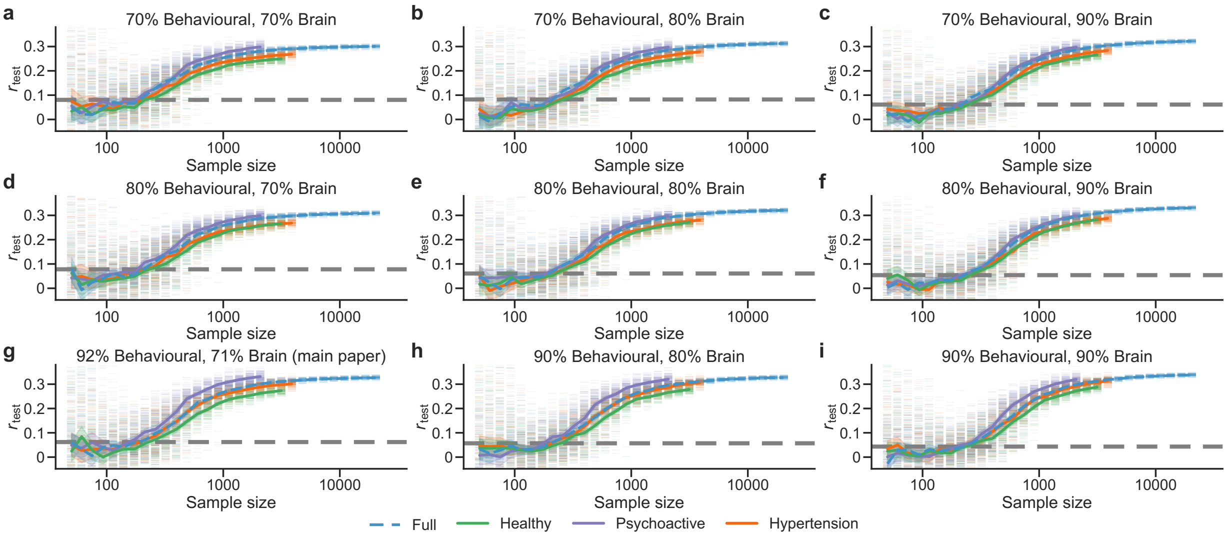


**Supplementary Figure 3: Replicability of canonical correlations across sample sizes and by cohort, with different numbers of principal components (CCA model with cross-validation sampling)**.

**a–i**: Average test set correlation by sample size for the Full cohort and for each other cohort featured in this analysis. Grey dashed line represents the 99th percentile of the null model distribution for the Full cohort at its greatest sample size. Error bands denote 95% confidence intervals.

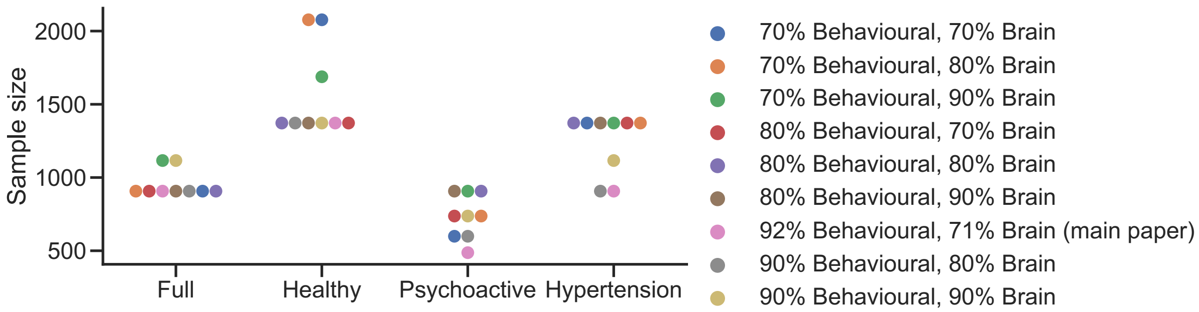


**Supplementary Figure 4: Smallest sample sizes needed to reach a mean test correlation value of at least 0.24 for each cohort when using different thresholds for dimensionality reduction (CCA model <u>with</u> cross-validation sampling)**.

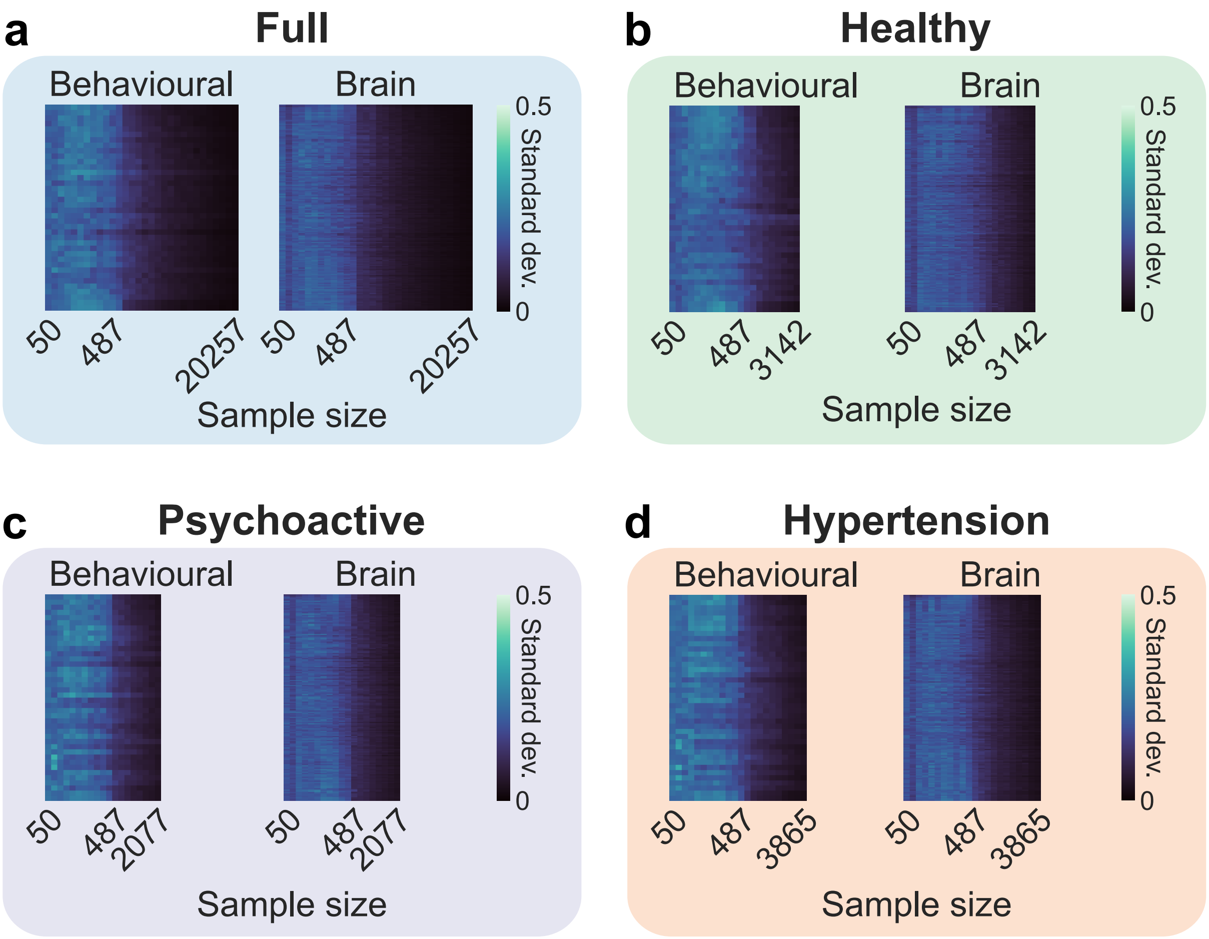


**Supplementary Figure 5: Standard deviation of loadings by sample size for brain and behavioural variables for each cohort (CCA model with cross-validation sampling).** Variable order is based on loading strengths at the largest sample size for each cohort. **a**: Full cohort. **b**: Healthy cohort. **c**: Psychoactive cohort. **d**: Hypertension cohort.

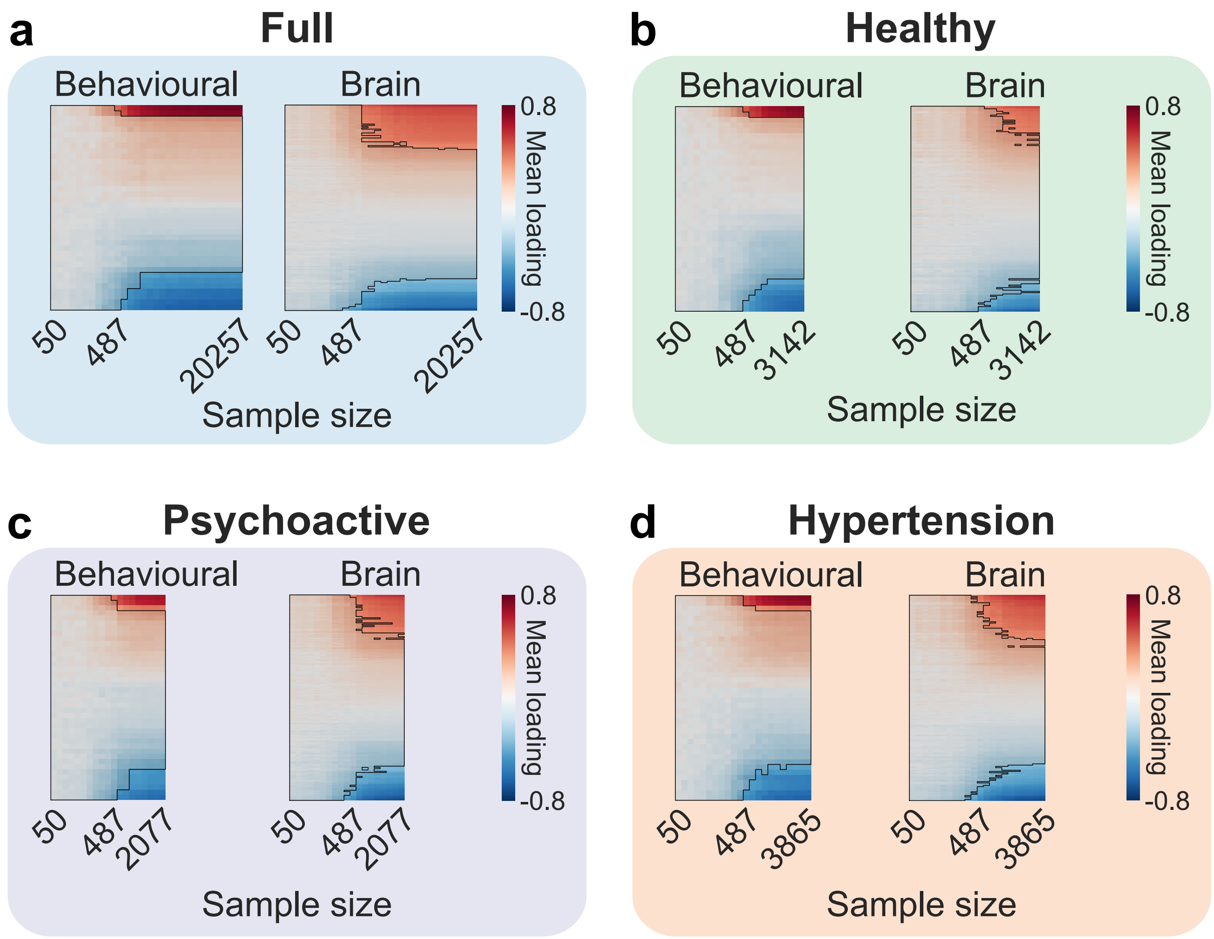


**Supplementary Figure 6: Mean loadings by sample size for brain and behavioural variables for each cohort (CCA model <u>without</u> cross-validation sampling).**
Variable order is based on loading strengths at the largest sample size for each cohort. The saturation indicates the strength of the loading. Loadings whose mean value falls within the 2.5–97.5 null model percentile range are masked. **a**: Full cohort. **b**: Healthy cohort. **c**: Psychoactive cohort. **d**: Hypertension cohort.

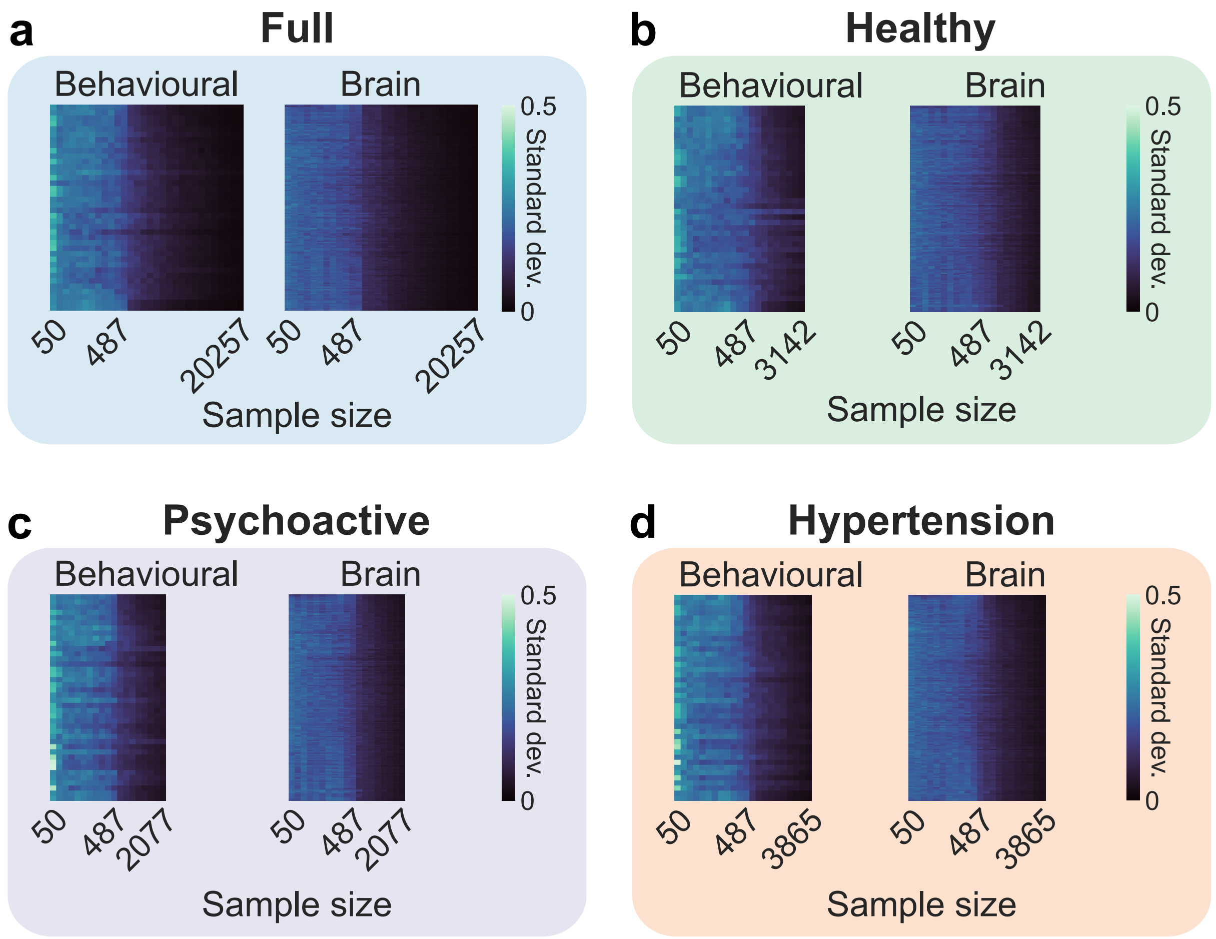


**Supplementary Figure 7: Standard deviation of loadings by sample size for brain and behavioural variables for each cohort (CCA model without cross-validation sampling).** Variable order is based on loading strengths at the largest sample size for each cohort. **a**: Full cohort. **b**: Healthy cohort. **c**: Psychoactive cohort. **d**: Hypertension cohort.

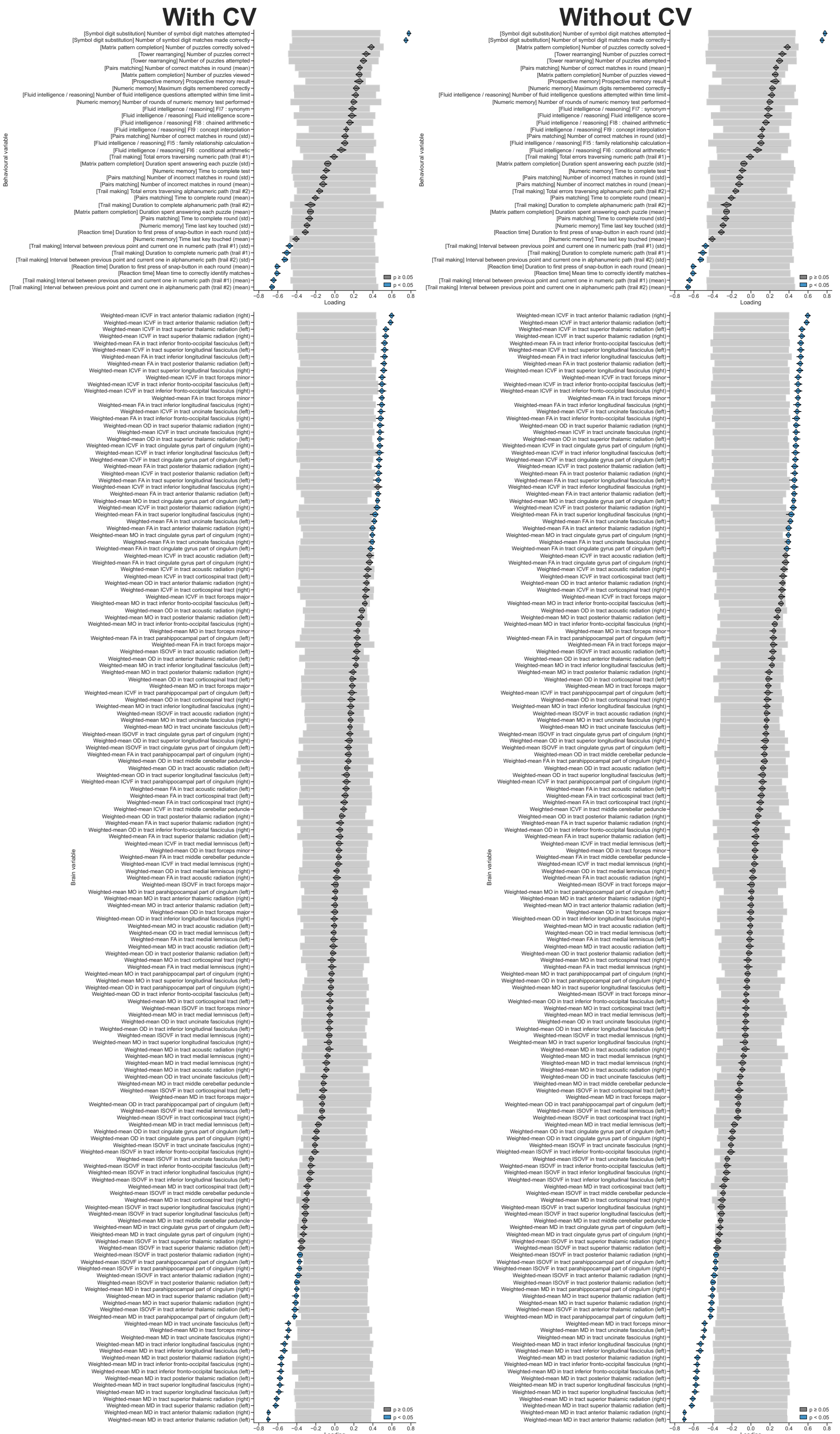


**Supplementary Figure 8: Complete loading distributions for Full cohort (CCA models <u>with</u> and <u>without</u> cross-validation sampling).**
Non-grey distributions denote variables that passed 2.5–97.5 null model percentile masking.

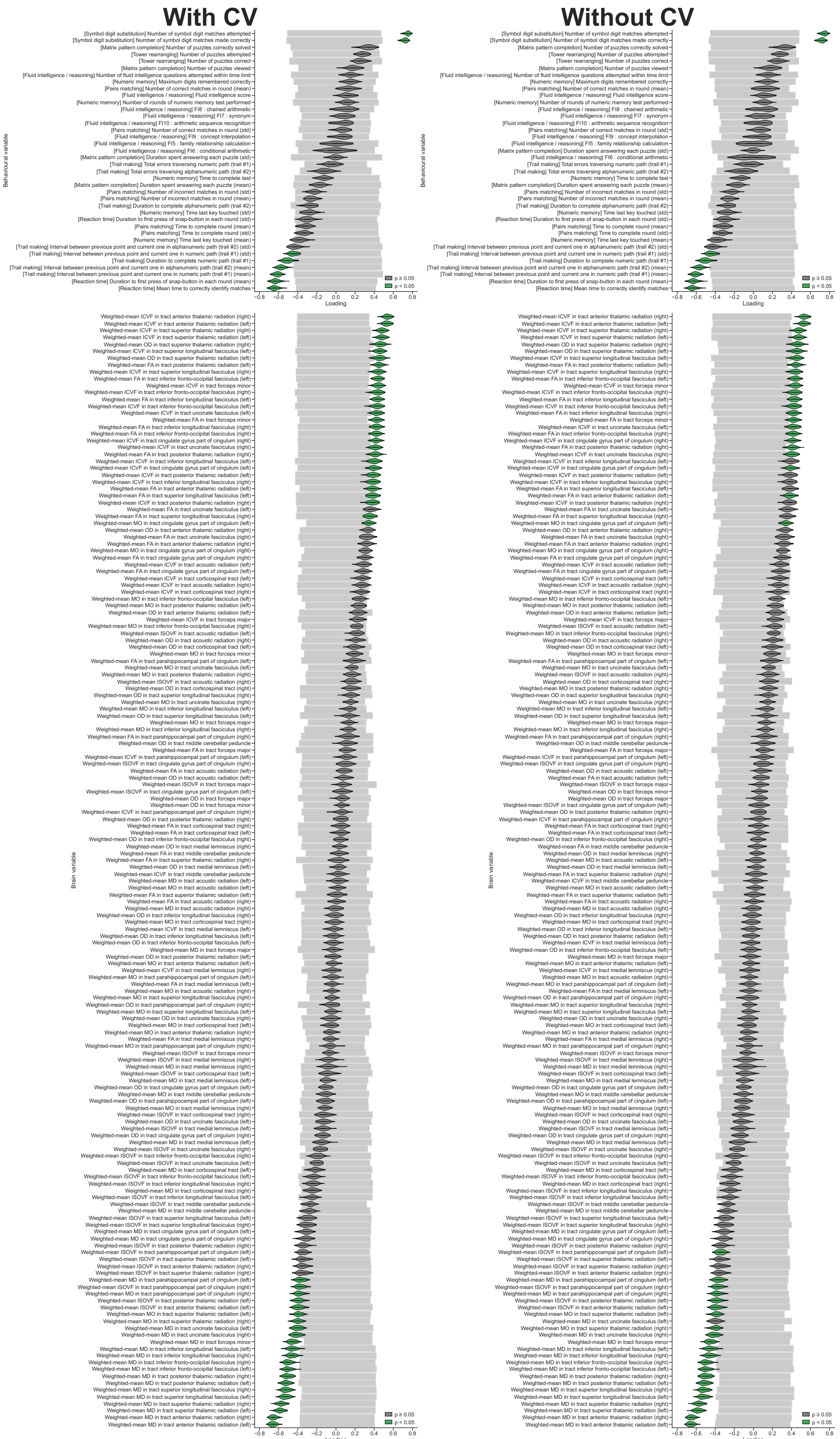


**Supplementary Figure 9: Complete loading distributions for Healthy cohort (CCA models <u>with</u> and <u>without</u> cross-validation sampling).**
Non-grey distributions denote variables that passed 2.5–97.5 null model percentile masking.

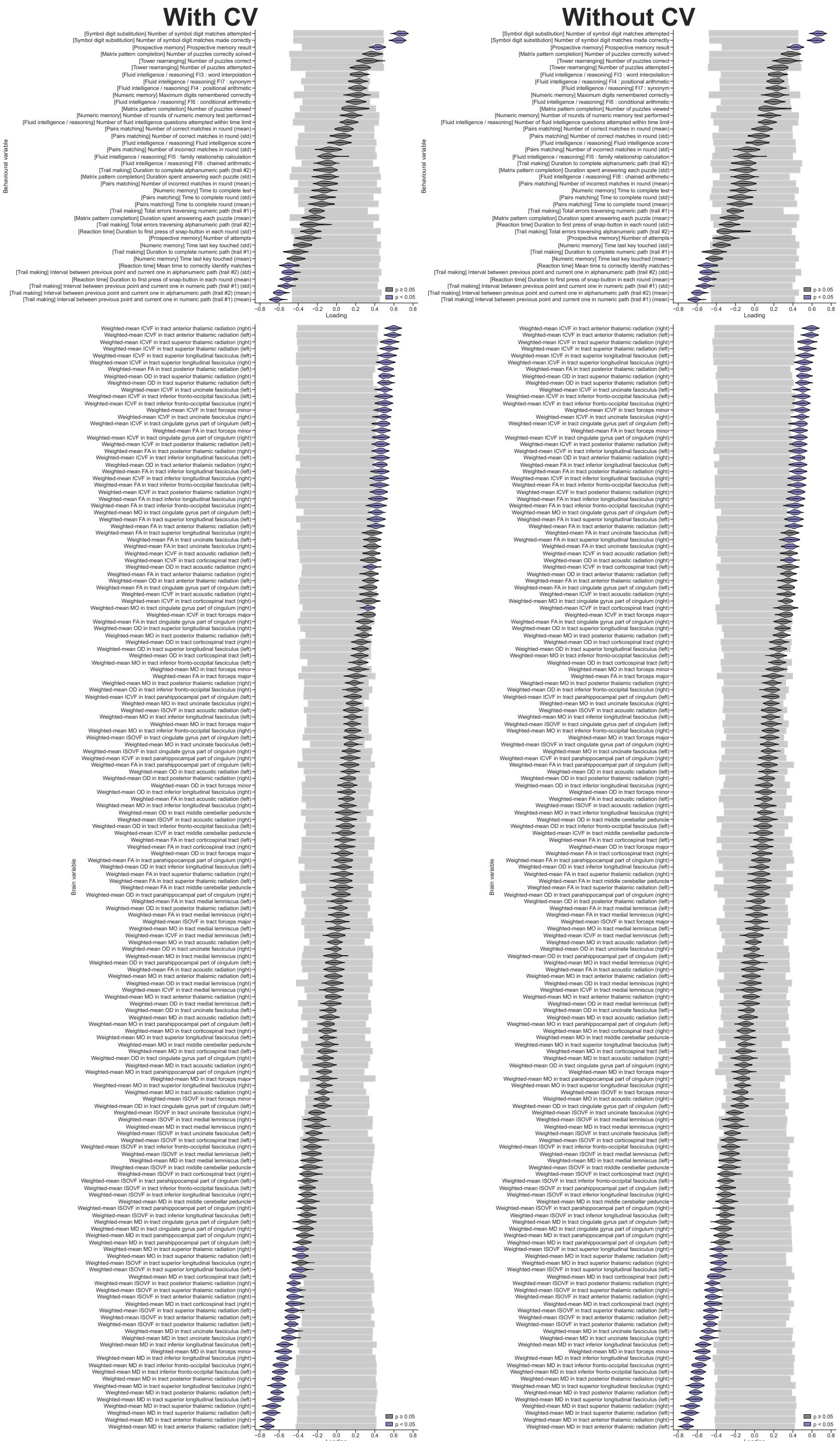


**Supplementary Figure 10: Complete loading distributions for Psychoactive cohort (CCA models <u>with</u> and <u>without</u> cross-validation sampling).**
Non-grey distributions denote variables that passed 2.5–97.5 null model percentile masking.

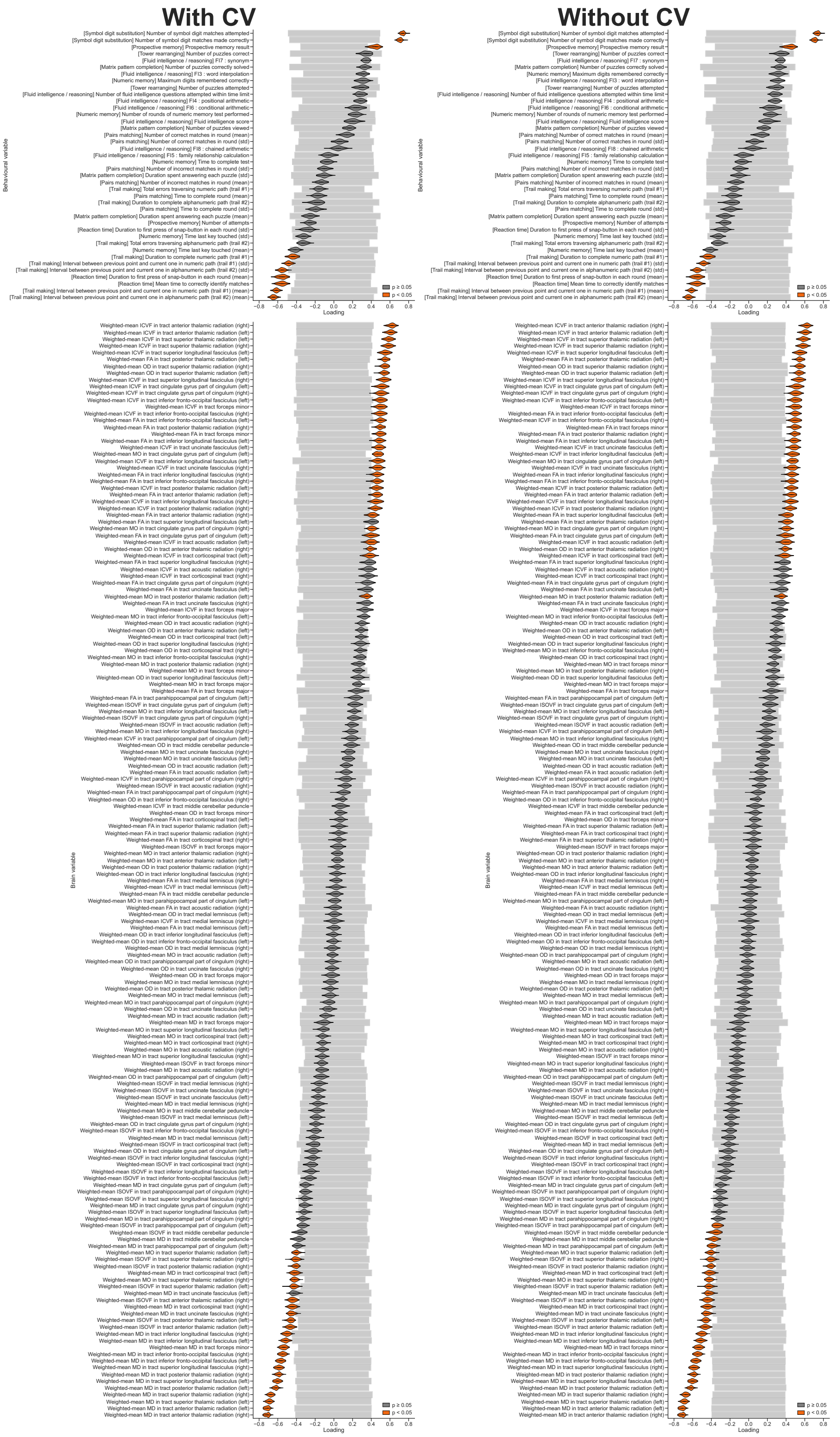


**Supplementary Figure 11: Complete loading distributions for Hypertension cohort (CCA models <u>with</u> and <u>without</u> cross-validation sampling).**
Non-grey distributions denote variables that passed 2.5–97.5 null model percentile masking.

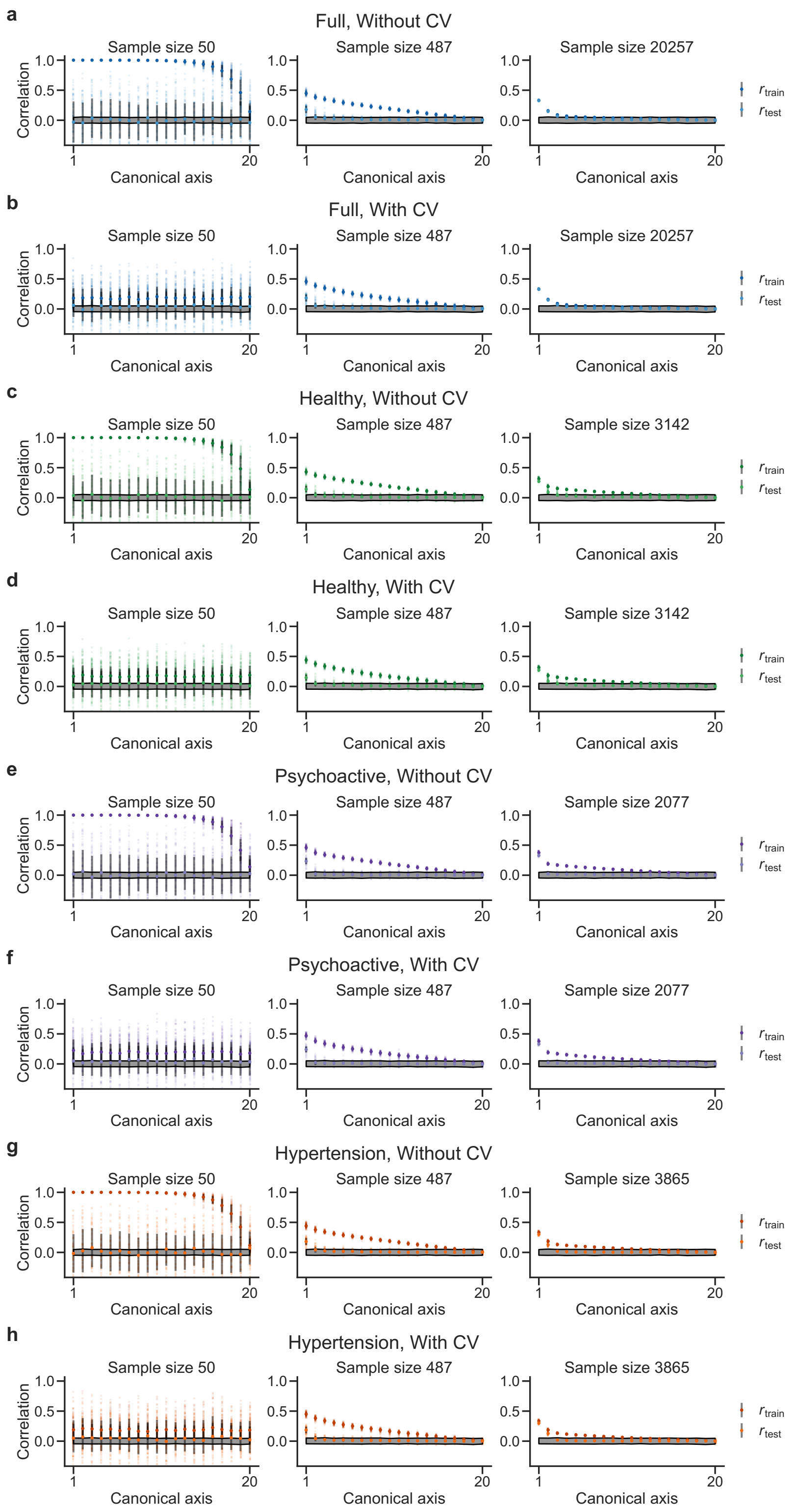


**Supplementary Figure 12: Canonical correlations for all axes.**
**a–h**: Mean train and test correlations for each canonical axis, cohort and CCA model type for three different sample sizes (50, 487, and maximum). Error bars indicate standard deviations.

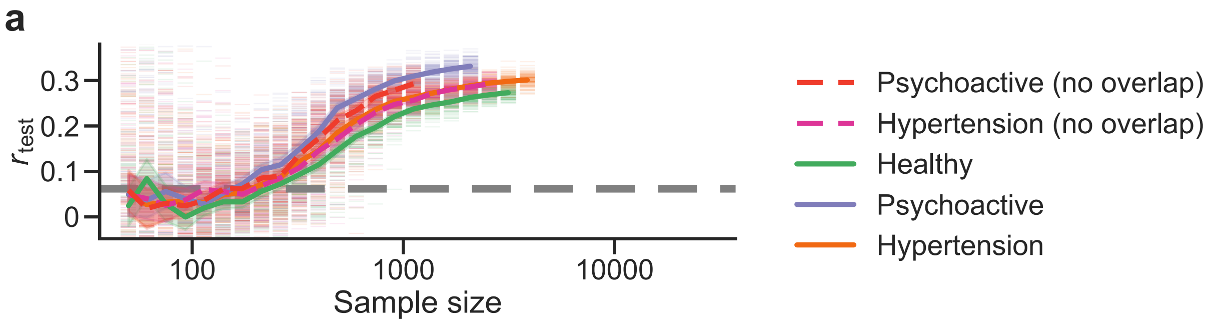


**Supplementary Figure 13: Replicability of canonical correlations across sample sizes and by cohort, with additional non-overlapping Psychoactive and Hypertension cohorts (CCA model with cross-validation sampling)**.
Average test set correlation by sample size for the Healthy, Psychoactive, Hypertension cohorts from the main analysis as well as non-overlapping subsets of the Psychoactive and Hypertension cohorts. Grey dashed line represents the 99th percentile of the null model distribution for the Full cohort at its greatest sample size. Error bands denote 95% confidence intervals.
